# Knockout of the tomato *HAIRY MERISTEM 4* alters phloem-characteristics and impairs development

**DOI:** 10.1101/2024.08.02.606343

**Authors:** Jackson Khedia, Abhay Pratap Vishwakarma, Ortal Galsurker, Shira Corem, Suresh Kumar Gupta, Tzahi Arazi

## Abstract

The HAIRY MERISTEM (HAM) gene family encodes Type I and II GRAS domain transcription factors in plants. Type II HAMs, predominantly expressed in meristems and regulated by microRNA171, are essential for maintaining undifferentiated meristems, a role conserved across various species. Conversely, the functions of Type I HAMs have been less characterized. In this study, we investigated the role of SlHAM4, a Type I HAM in tomato. Using publicly available expression data and a GUS reporter gene driven by the native *SlHAM4* promoter, we determined that *SlHAM4* is predominantly expressed in phloem tissues. CRISPR-induced *SlHAM4* loss-of-function mutations (*slham4^CR^*) resulted in a range of shoot and fruit abnormalities, which were fully reversed by reintroducing *SlHAM4* under its native promoter in the mutant background. Mutant abnormalities included increased anthocyanin pigmentation in the leaf and sepal primordia, reminiscent of the phenotypes observed in certain Arabidopsis mutants with compromised phloem, and development of simpler leaves, which was associated with reduction in external phloem area in the leaf rachis. In addition, *slham4^CR^* plants produced significantly smaller fruits of which a fraction of them exhibited catface-like scars, attributed to tears which occurred in the pericarp of mutant ovaries following fruit set. Transcriptome analysis of the wild-type looking mutant ovaries at anthesis revealed specific downregulation of genes implicated in phloem development and functions, in particular those expressed in companion cells (CC). We propose that SlHAM4 is necessary for proper phloem function in part by regulating the expression of a suite of CCs genes that encode essential phloem proteins.

**One-sentence summary:** *SlHAM4* is predominantly expressed in the phloem and its knockout alters phloem- characteristics and impair development highlighting its requirement for proper phloem functionality.

## Introduction

The plant HAIRY MERISTEM (HAM) gene family, initially identified in Petunia (Stuurman *et al*., 2002), encodes GRAS domain transcription factors. In Arabidopsis, four HAM homologs (HAM1-HAM4) have been identified and are categorized into two groups, Type I and Type II, according to phylogenetic analysis (Engstrom *et al*., 2010). The Petunia *ham* mutant exhibits hairy shoot apical meristem (SAM) due to its premature differentiation, as well as termination of lateral organ development (Stuurman *et al*., 2002).

In Arabidopsis, the Type II HAM group, encompassing HAM1, HAM2, and HAM3 (alternatively known as LOST MERISTEM1 (LOM1), LOM2, and LOM3), exhibits primary expression in shoot and root meristems, as well as in the provascular tissues (Engstrom *et al*., 2010; Zhou *et al*., 2015a). These genes are subject to negative regulation by microRNA171 (miR171), which mediates transcript cleavage, thereby modulating gene expression (Llave *et al*., 2002). Individual loss-of-function mutations in these genes typically do not yield substantial developmental anomalies. However, the phenotypic consequences of combined mutations or miR171 overexpression are significant, including disorganized SAM and axillary meristem architecture that are associated with early termination of SAM and a reduction in axillary shoot branching (Engstrom *et al*., 2010; Zhou *et al*., 2015a). Additional analysis of mutants has elucidated that HAM1 and HAM2 predominantly contribute to the maintenance of undifferentiated SAMs and the initiation of new axillary stem cell niches. Conversely, HAM3 appears to play a comparatively minor role in the maintenance of shoot stem cells primarily involved in the development of axillary meristems (Han *et al*., 2020). Notably, HAM1 and HAM2 directly interact with the WUSCHEL (WUS) protein, functioning as transcriptional co-factors. In addition, they prevent *CLAVATA3* (*CLV*3) expression within the inner cells of the SAM. These activities are pivotal in controlling shoot stem cell production and maintaining the indeterminacy of the SAM. In several flowering plants, the HAM gene family maintains indeterminate SAMs and promotes new axillary meristem formation, suggesting that this function is conserved across several plant species (Geng and Zhou, 2021).

Type I HAM genes show variation in their miR171 target sequences, unlike Type II HAM genes. Some Type I HAM genes retain the conserved miR171 binding sequence, while others, such as HAM4 in Arabidopsis, have lost it (Engstrom *et al*., 2010; Llave *et al*., 2002), suggesting potential functional diversification. The expression of the Arabidopsis Type I gene *HAM4* (also referred to as *SCARECROW- LIKE15*) is restricted to the vasculature of leaves, roots, stems, and siliques, particularly in the phloem system, including companion cells (Gao *et al*., 2015; Zhou *et al*., 2015a). HAM4 has been shown to physically interact with WUSCHEL-RELATED HOMEOBOX4 (WOX4), which is expressed in procambial cells that define the vascular stem cell niche. This interaction, coupled with their coinciding expression patterns, suggest a potential role for HAM4 as a cofactor of WOX4 (Zhou *et al*., 2015a). However, phenotypic analysis of *scl15/ham4* mutants revealed no significant abnormalities in plant growth, except that they were slightly smaller and exhibited a minor delay in flowering compared to wild-type plants. (Gao *et al*., 2015; Zhou *et al*., 2015a). In addition, HAM4/SCL15 has been identified as a partner of HISTONE DEACETYLASE19 (HDA19) in Arabidopsis seedlings. Microarray analysis of mutant *scl15/ham4* seedlings revealed that loss of *HAM4/SCL15* led to significant changes in gene expression, with numerous seed maturation genes being upregulated in vegetative tissues. The upregulation of a fraction of them was associated with increased histone acetylation, an epigenetic mark associated with expression activation. These results suggest that SCL15 acts as an HDA19-interacting regulator, repressing a subset of seed maturation genes through histone deacetylation (Gao *et al*., 2015). Nevertheless, whether HAM4 plays specific roles in phloem development and function remains largely elusive.

Tomato contains three HAM homologs—the Type II SlHAM and SlHAM2, and the Type I SlHAM4 (Solyc02g085600). *SlHAM* and *SlHAM2* are targeted for cleavage by miR171 and are abundant in the shoot and floral meristems, as well as in compound leaf primordia. Silencing *SlHAM* and *SlHAM2* leads to over proliferation of stem cells in the periphery of the meristems and in the organogenic compound leaf rachis, when specifically silenced in leaves, suggesting that SlHAM and SlHAM2 play conserved roles in meristem maintenance (Hendelman *et al*., 2016). Unlike *SlHAM* and *SlHAM2*, *SlHAM4* is not targeted by miR171 and is practically absent from meristems and compound leaf primordia (Hendelman *et al*., 2016), suggesting that it is not playing a major role in meristem maintenance and may have other functions.

In this study, we explored the functions of *SlHAM4* in tomato. Our findings reveal that SlHAM4 is predominantly expressed in phloem tissues. Loss-of-function mutations in *SlHAM4* induced by CRISPR/Cas9 (*slham4^CR^*), resulted in various shoot and fruit abnormalities. These included increased anthocyanin pigmentation, simpler leaves, and smaller fruits with catface-like scars. These defects were fully reversed by reintroducing *SlHAM4* under its native promoter and were associated with reduction in external phloem area in mutant leaves rachises and downregulation of anthesis ovary genes involved in phloem development and function, particularly those expressed in companion cells (CC). Altogether, our results suggest that SlHAM4 acts to regulate the expression of phloem-associated genes which are essential for phloem function and hence tomato development.

## Results

### The *slham4^CR^* mutations are associated with abnormal shoot and fruit development

To functionally characterize *SlHAM4*, we first employed CRISPR/Cas9 technology to knockout its gene (Fig. 1A). This strategy yielded two distinct *SlHAM4* mutant alleles with deletions of 3 and 4 base pairs, subsequently named *slham4^CRΔ3^* and *slham4^CRΔ4^*, respectively (Fig. 1B). The SlHAM4 protein is a member of the GAI, RGA, and SCR (GRAS) family of transcriptional factors (Bolle, 2004). It is characterized by a variable N-terminus and a highly conserved GRAS domain, which includes leucine-rich regions (LRI and LRII) surrounding a VHIID motif (V^280^HVID^284^) and a C-terminal SAW motif (S^532^AW^534)^ (Fig. 1C). The *slham4^CRΔ3^* mutation likely leads to the loss of the conserved Leu^271^ within the LRI region (Fig. 1C and D). The *slham4^CRΔ4^* mutation introduces a premature stop codon, resulting in a significantly truncated SlHAM4 protein missing a substantial portion of its GRAS domain (Fig. 1D), suggesting that it represents a loss-of-function mutation.

**Figure 1.**
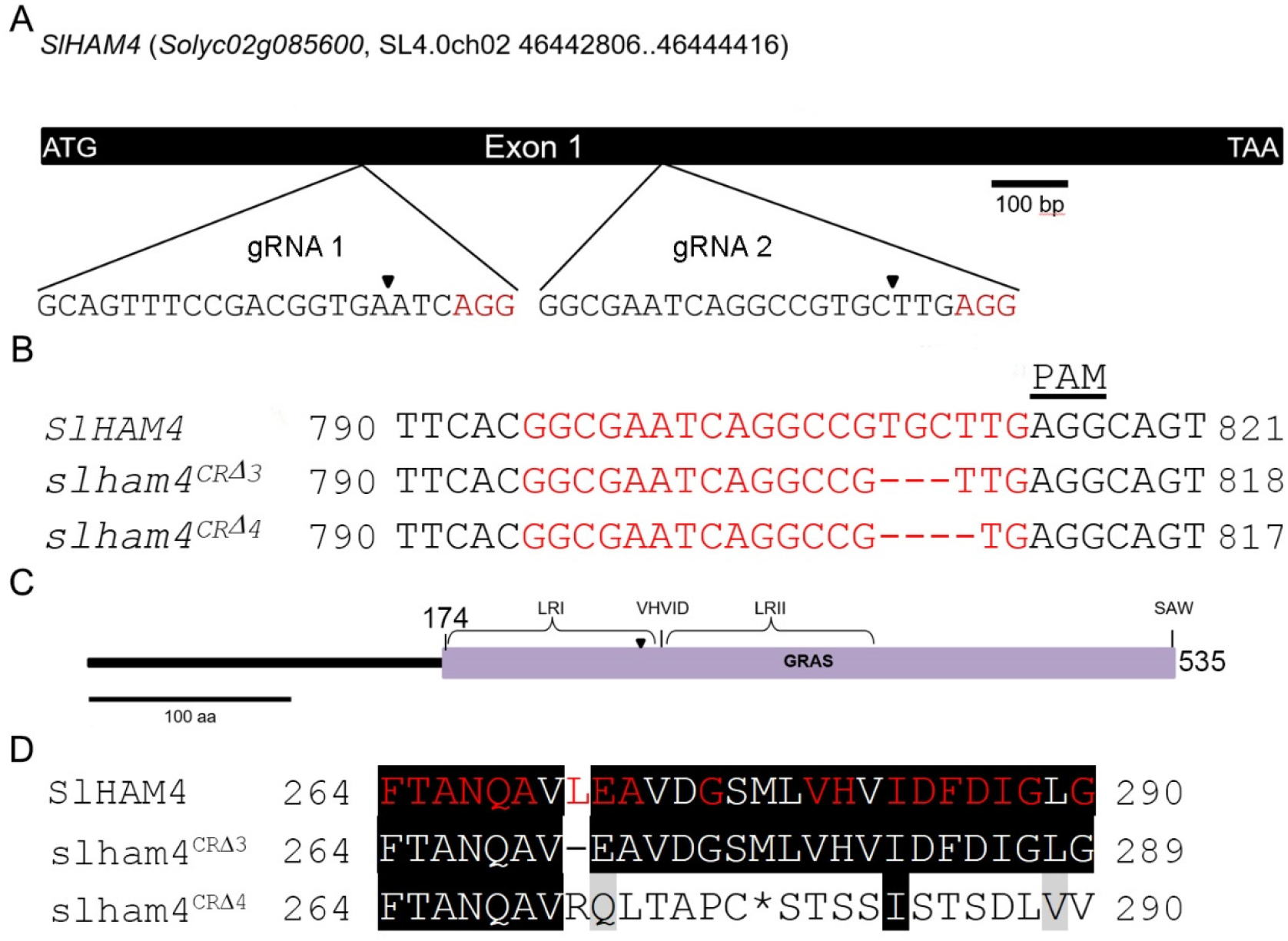
CRISPR/Cas9-mediated mutagenesis of the *SlHAM4* gene. (A) Schematic of the *SlHAM4* gene structure with guide RNA (gRNA) target sites marked. Expanded views show gRNA target sequences with PAM motifs (red); arrowheads indicate expected Cas9 cleavage points. (B) Alignment of CRISPR mutant alleles (*slham4^CR^*) with the wild-type *SlHAM4* sequence. gRNA target sequences are highlighted in red, and the PAM motif is noted. Numbering is from the start codon. (C) Representation of SlHAM4 protein architecture. The GRAS domain, key sequence motifs, and regions are shown. The conserved Leu^271^ location is marked with an arrowhead. (D) Alignment of mutant and wild-type SlHAM4 protein sequences. Amino acids in the GRAS domain conserved across species are highlighted in red, according to the NCBI Conserved Domain Database (https://www.ncbi.nlm.nih.gov/cdd/). Numbering starts from the start codon, with an asterisk (*) indicating a premature stop codon.

To investigate the role of *SlHAM4* in tomato development, we compared the vegetative and reproductive phenotypes of the *slham4^CR^*mutants to the M82 parental line (henceforth will be named wild type). Initially, both *slham4^CRΔ3^* and *slham4^CRΔ4^* mutants exhibited normal germination and seedling growth, mirroring the phenotype of wild-type seedlings (Fig. 2A). However, as they matured the mutants could be distinguished from wild-type plants based on the reduced density of their shoots, attributed to the mutants’ compound leaves bearing fewer leaflets (Fig. 2B and C). Furthermore, the *slham4^CR^* mutants displayed pronounced purple pigmentation in leaf primordia and flower bud sepals, a sign of anthocyanin accumulation, a phenotype never observed in wild-type plants under identical growth conditions (Fig. 2D). The purple pigmentation was also present in the sepals of *slham4^CR^* flowers at anthesis, although no other differences from the wild-type flowers were observed (Fig. 2E and F). Unfertilized wild type tomato flowers senesce around 5 days after anthesis, marked by petal dehydration and weak chlorosis of sepals. Interestingly, while petal dehydration occurred in senescing *slham4^CR^* mutant flowers, their sepals exhibited strong chlorosis and browning of certain areas at their base, deviating from the typical sepal senescence pattern observed in wild type (Fig. 2G).

**Figure 2.**
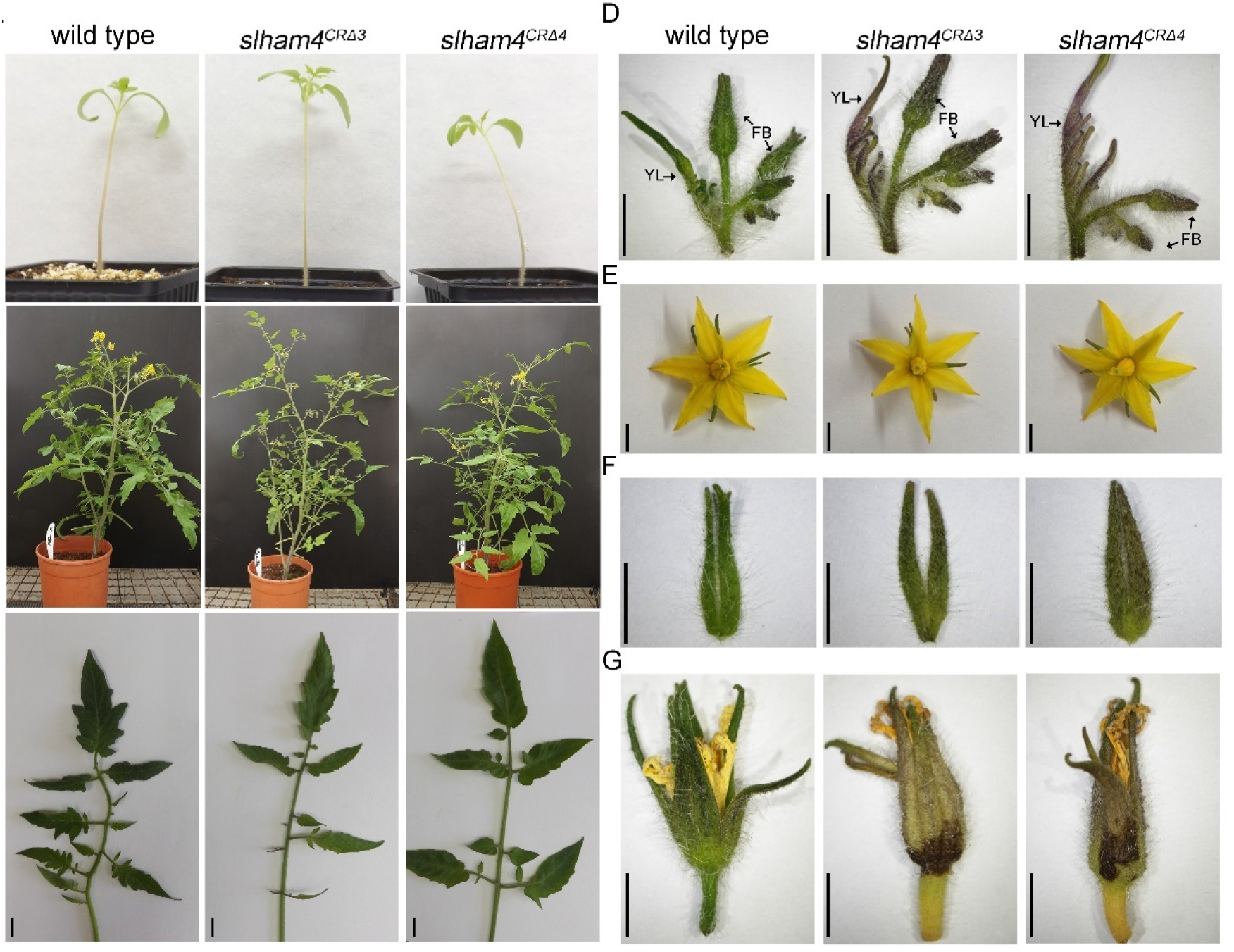
Comparison of M82 (wild type) and *slham4^CR^* shoot and flower phenotypes. (A) Seedlings at 14 days post-germination. (B) Mature plants cultivated in standard greenhouse conditions. (C) A representative fully expanded leaf. (D) Representative young leaves and inflorescences at the sympodial shoot. YL, young leaf; FB, flower bud. Note the increased purple pigmentation of mutant young leaves and sepals. (E) Anthesis flowers. (F) Detached sepals from anthesis flowers. (G) Representative flowers not subjected to fertilization, displaying senescence; mutant sepals exhibit noticeable dark brown areas. Scale bars = 1 cm (C) and 5 mm (D-G).

At anthesis, the *slham4^CR^*flower pistils appeared morphologically identical to wild type controls (Fig. 3A). Consistent with this, and despite relatively high *SlHAM4* expression in the ovary placenta and columella (Fig. S1A to C), detailed histological analysis of both wild-type and mutant ovaries at anthesis revealed no clear differences in their overall structure and vasculature arrangement (Fig. 3E). However, we noticed that following fruit set, a fraction of *slham4^CR^* young immature green fruits exhibited small tears in their pericarp (Fig. 3B). As the fruit grew, these tears expanded, eventually exposing the locules to air resulting in scarring (Fig. 3C and D). Histological analysis of *slham4^CRΔ4^* young immature green fruits revealed indentations in the pericarp containing collapsed cells (Fig. 3F). We hypothesized that the damaged tissue’s inability to grow with the rest of the pericarp resulted in a tear that expanded as the fruit developed. To test this, we created a ∼1 mm indentation in the pericarp of M82 anthesis ovaries by gently crushing it with the tip of a pencil. This manipulation led to the formation of a tear that expanded as the fruit grew (Fig. 3G), suggesting that localized pericarp damage can lead to significant fruit scars. The observed fruit scars in *slham4^CR^*mutants varied in size and shape (Fig. 3C and Fig. S2A to C). This fruit pericarp scarring phenotype is highly reminiscent of the tomato fruit catfacing syndrome (Peet, 2009; Knavel and Mohr, 1969). Notably, catfacing occurred in many, but not all, *slham4^CRΔ4^* fruits, suggesting incomplete penetrance. Under our greenhouse conditions, 30-70% of *slham4^CRΔ4^* fruits displayed catfacing, while both heterozygous *slham4^CRΔ4(-/+)^* and wild- type fruits were completely devoid of catfacing (Fig 3H). Mutant fruits with and without catfacing ripen normally, and their internal morphology is not significantly different from that of wild-type tomatoes (Fig. 3D and E), except that their final size is significantly smaller than wild-type fruits (Fig. 3I).

**Figure 3.**
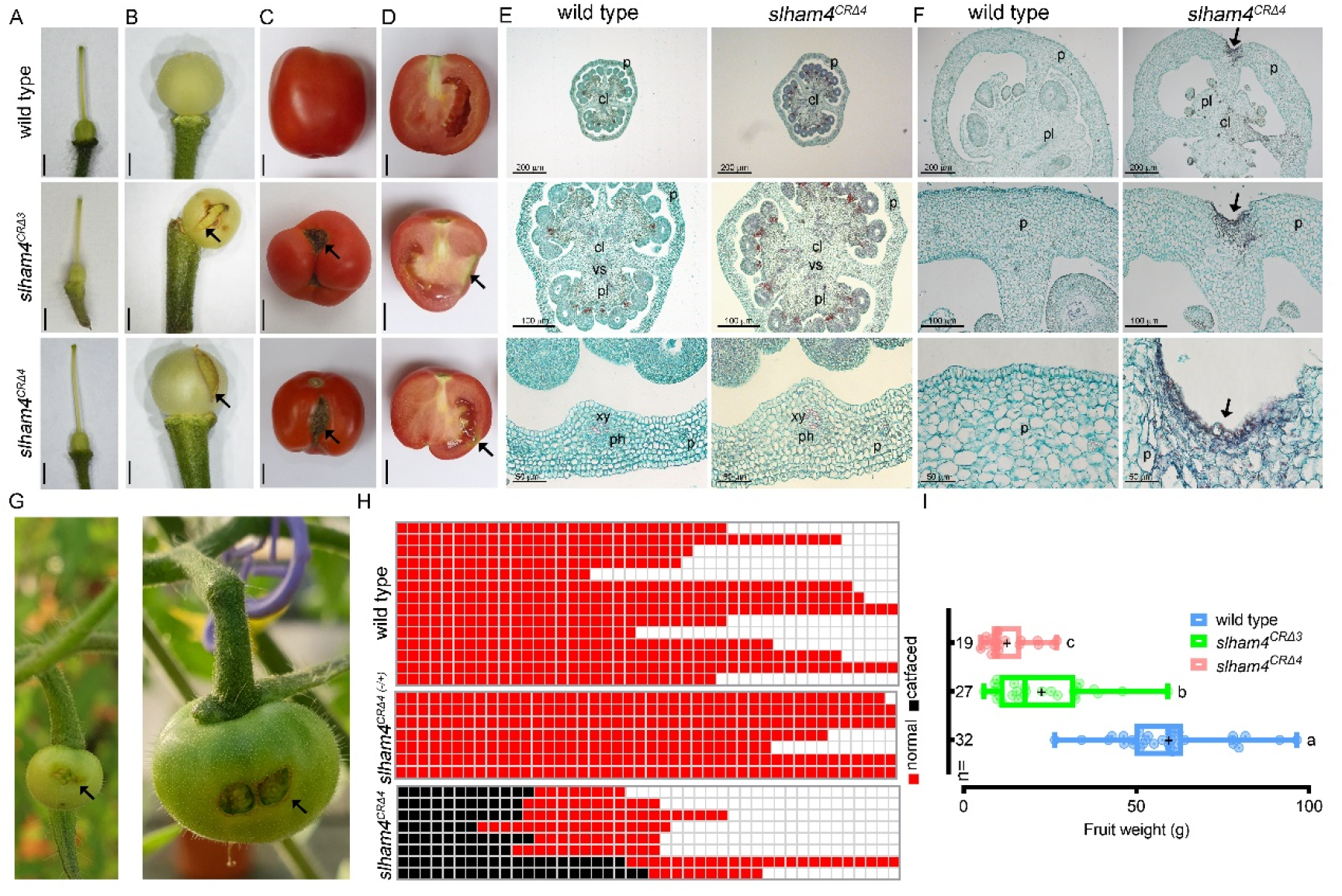
Characterization of M82 (wild type) and *slham4^CR^* fruits. (A) Isolated pistils from anthesis flowers. (B) Young immature green fruits. Arrows mark the pericarp ruptures in mutants. (C and D) Red ripe fruits and their longitudinal sections, respectively. Arrows point to catface-like scars. Scale bars = 2 mm (A and B) and 1 cm (C and D). (E and F) Cross-sections of anthesis ovaries and 4 mm stage immature green fruits, respectively, stained with safranin and fast-green, where the red staining delineates xylem vasculature. The middle and bottom panels show close up views of respective upper panel. P, pericarp; cl, columella; pl, placenta; vs, vascular tissue; xy, xylem; ph, phloem. In F, arrows mark the indentation in the mutant fruit pericarp. (G) Young immature and immature green M82 fruits developed from anthesis ovaries with manually crushed pericarp, resulting in a ∼1 mm indentation. The resulting catface-like scars are indicated by arrows. (H) Diagram illustrating the prevalence of catface formation in fruits from wild type, heterozygous (*slham4^CRΔ4-/+^*), and homozygous (*slham4^CRΔ4^*) plants, with each row representing an individual plant. Plants were cultivated in greenhouse nested plots. (I) Average fruit weight of indicated genotypes. The median and average are indicated by a line and +, respectively; n = number of fruits; Different letters indicate significance (*P* < 0.01) as determined by Tukey–Kramer multiple comparison test.

### *SlHAM4* transgenic expression reverts mutant phenotypes and delineates its promoter activity

To confirm the specific contribution of *SlHAM4* to the observed *slham4^CR^* phenotypes, we investigated whether reintroducing a functional copy could counteract the effects of the loss-of-function mutation. First, we generated transgenic M82 plants expressing *SlHAM4* driven by the constitutive cauliflower mosaic virus *35S* (*35S*) promoter (Fig. S3A). Among the regenerated lines, *35S::SlHAM4-5* displayed the highest *SlHAM4* overexpression (Fig. S3B) with no adverse effects on plant development (Fig. S3C and Fig 4A and B). Crossing this line with homozygous *slham4^CRΔ4^* mutant and analyzing the *35S::SlHAM4-5 slham4^CRΔ4^* F2 progeny revealed no difference in flower senescence and fruit development compared to wild type (Fig. 4A-C). The elimination of fruit catfacing coincided with a six-fold increase in *SlHAM4* expression in *35S::SlHAM4-5 slham4^CRΔ4^* anthesis ovaries compared to wild type (Fig. 4D).

**Figure 4.**
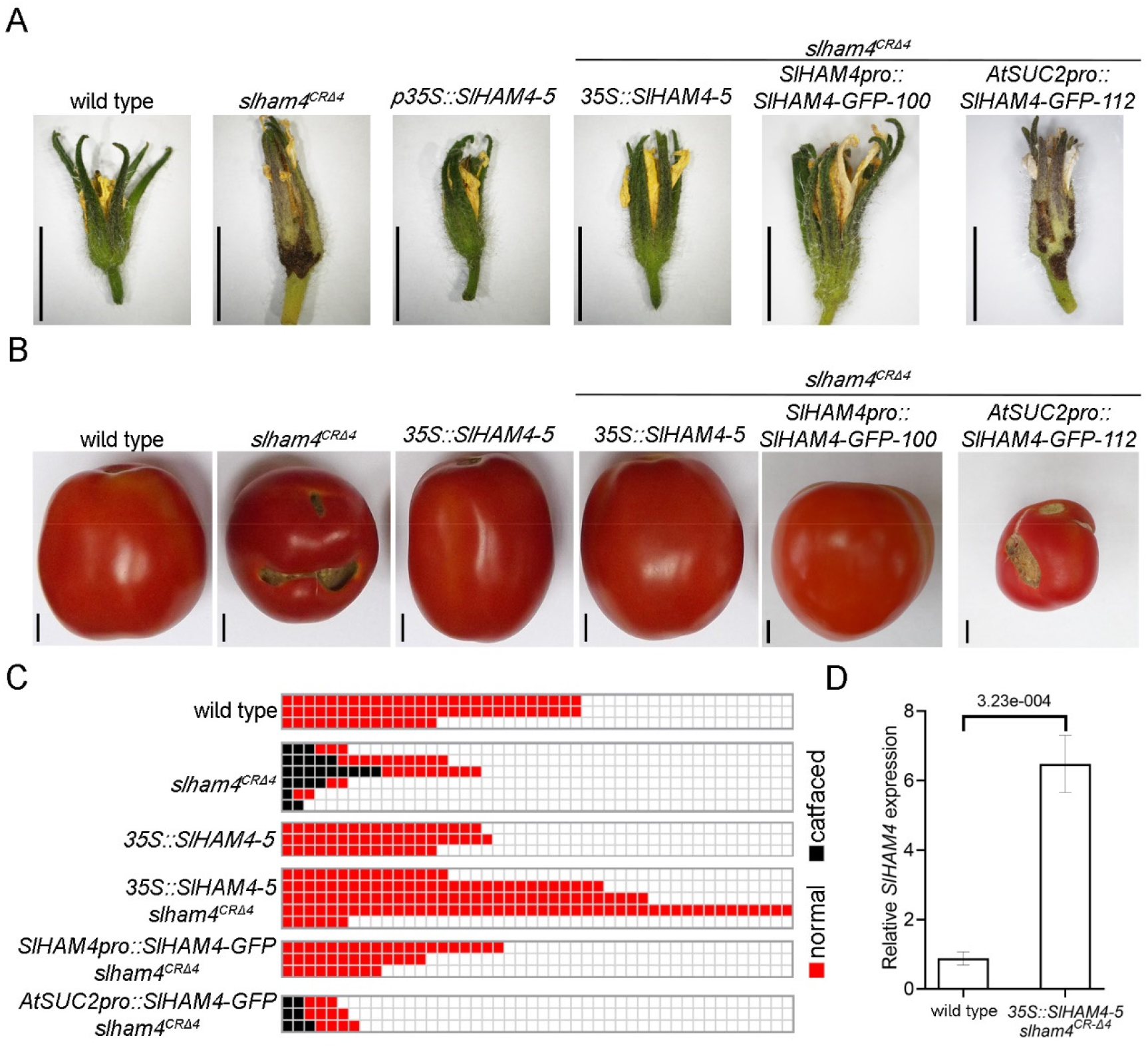
Functional complementation of *slham4^CRΔ4^* mutation. (A and B) Images of representative senescing flowers (A) and red ripe fruits (B) from M82 (wild type) and indicated genotypes. Scale bars = 1 cm. (C) Schematic representation of catface development in fruits of indicated genotypes. Each column represents a single plant. (D) Quantitation of *SlHAM4* transcript levels in M82 (wild type) and *35S*::*SlHAM4-5 slham4^CRΔ4^* anthesis ovaries, normalized to *SlTIP41* as the reference gene. Error bars indicate ±SD over 3 biological replicates. The P value as determined by students t-test is shown.

The *SlHAM4* gene is situated on chromosome two, located 8.82 Kb upstream of *Solyc02g085590* and 0.48 Kb downstream of *Solyc02g085610* (Fig. S3D). To delineate the *SlHAM4* endogenous promoter we generated transgenic *slham4^CRΔ4^* plants expressing a SlHAM4-GFP fusion protein driven by a 4 kb fragment of the putative promoter located upstream of *SlHAM4* start codon (Fig. S3D and E). Among regenerated transgenic lines, *SlHAM4pro::SlHAM4-GFP slham4^CRΔ4^* lines 8, 100 and 102 regained normal flower senescence and were completely devoid of fruit catfacing, suggesting sufficient activity of the 4Kb promoter fragment (Fig. 4A-C). Collectively, these experiments strongly support a direct link between SlHAM4 function and the observed phenotypes in *slham4^CR^* mutants. Moreover, the similar phenotypic abnormalities between *slham4^CRΔ3^* and *slham4^CRΔ4^* mutants (Fig. 2 and Fig. 3) reinforce the classification of *slham4^CRΔ3^* as a loss-of-function allele, highlighting the importance of the conserved Leu^271^ residue in the GRAS domain for the proper function of SlHAM4.

### *SlHAM4* is predominantly expressed in the phloem

To explore the tissue specific expression pattern of *SlHAM4*, we initially quarried public tomato databases for *SlHAM4* expression. These revealed that in seedlings, *SlHAM4* is predominantly expressed in the shoot relative to the roots (Fig. S1A inset). In mature plants, SlHAM4 is expressed in both vegetative and reproductive tissues, with the highest expression levels detected in anthesis flowers and orange fruit pericarp (Fig. S1A). In anthesis flower ovary, *SlHAM4* expression is strongest in the placenta and columella, tissues known for their rich vasculature (Gillaspy *et al*., 1993) (Fig. S1B and C). In developing and ripening fruits, *SlHAM4* is predominantly expressed in the pericarp vasculature and its levels increase progressively throughout fruit development and ripening, peaking in orange and red fruits (Fig. S1B to D).

To investigate the spatial distribution of *SlHAM4* in detail, we analyzed its promoter β-glucuronidase (GUS) activity in transgenic M82 tomato plants. We used a reporter gene construct, SlHAM4pro::GUS, which was generated by cloning the 4 kb fragment of the *SlHAM4* promoter, proven effective in driving native *SlHAM4* expression and restoring wild-type phenotypes in *slham4^CRΔ4^* (Fig. 4), upstream the GUS reporter gene (Fig. S4A). Analyzing 12 independent T0 transgenic plants, we observed vasculature-associated GUS staining in the leaves of 7 lines (Fig. S4B). These transgenic plants were allowed to set seeds and GUS activity was assayed in their transgenic progeny tissues. GUS staining was not observed in mature embryos extracted from 1 day after imbibition (DAI) transgenic seeds (Fig. 5A). At 1 d after germination (DAG), GUS staining was observed only in the region overlapping the transgenic cotyledons midveins (Fig. 5B). This GUS staining pattern was observed also in 3 DAG cotyledons and in addition, at that stage, weak non-specific GUS staining was observed in the hypocotyl (Fig. 5C). At 7 DAG, stronger GUS staining was observed in the cotyledons midvein, as well as in their minor veins. In addition, a weak but specific GUS staining was observed in the vasculature of the hypocotyl (Fig. 5D). At 18 DAG, GUS activity was found throughout the vascular tissues of the cotyledons, leaf primordia and stem (Fig. 5E). At 27 DAG, developing leaves showed GUS activity in their veins (Fig. 5F) and GUS activity was also observed in the stem vasculature (Fig. 5G). In line with the *SlHAM4* relatively weak expression in M82 seedling roots (Fig. S1A inset), until 27 DAGs, GUS staining was not detected in the roots (Fig. 5A to E). Histological sections of 27 DAG *SlHAM4pro::GUS* seedling stem (Fig. 5H and I) and cotyledon major and minor veins (Fig. S5A to D) revealed that GUS staining was confined to the phloem tissue within a vascular bundle. Within the phloem tissue, staining was detected in companion cells (CCs) and sieve elements (SEs) (Fig. 5I, Fig. S5B and C). As the SEs are enucleated (Imlau *et al*., 1999), the observed SEs staining likely reflects transport of the GUS enzyme or movement of the substrate cleavage product from their adjacent CCs. To explore the expression pattern of *SlHAM4* during fruit development, we characterized the GUS staining in ovaries at anthesis and in developing and ripening fruits. Dispersed but specific GUS staining was observed in the placenta and columella of ovaries at anthesis and in very young immature fruits (Fig. 5J, K and Fig. S4C). As the fruits matured and ripened, GUS staining intensity increased and became restricted to the vasculature (Fig. 5L, N and Fig. S4C). These patterns align with the native expression of *SlHAM4* (Fig. S1B to D), supporting its association with vasculature in fruits. Additionally, histology of GUS stained mature green fruit vasculature bundle revealed specific staining in phloem-associated cells (Fig. 5O and P), consistent with its spatial expression in the vascular bundles of seedlings.

**Figure 5.**
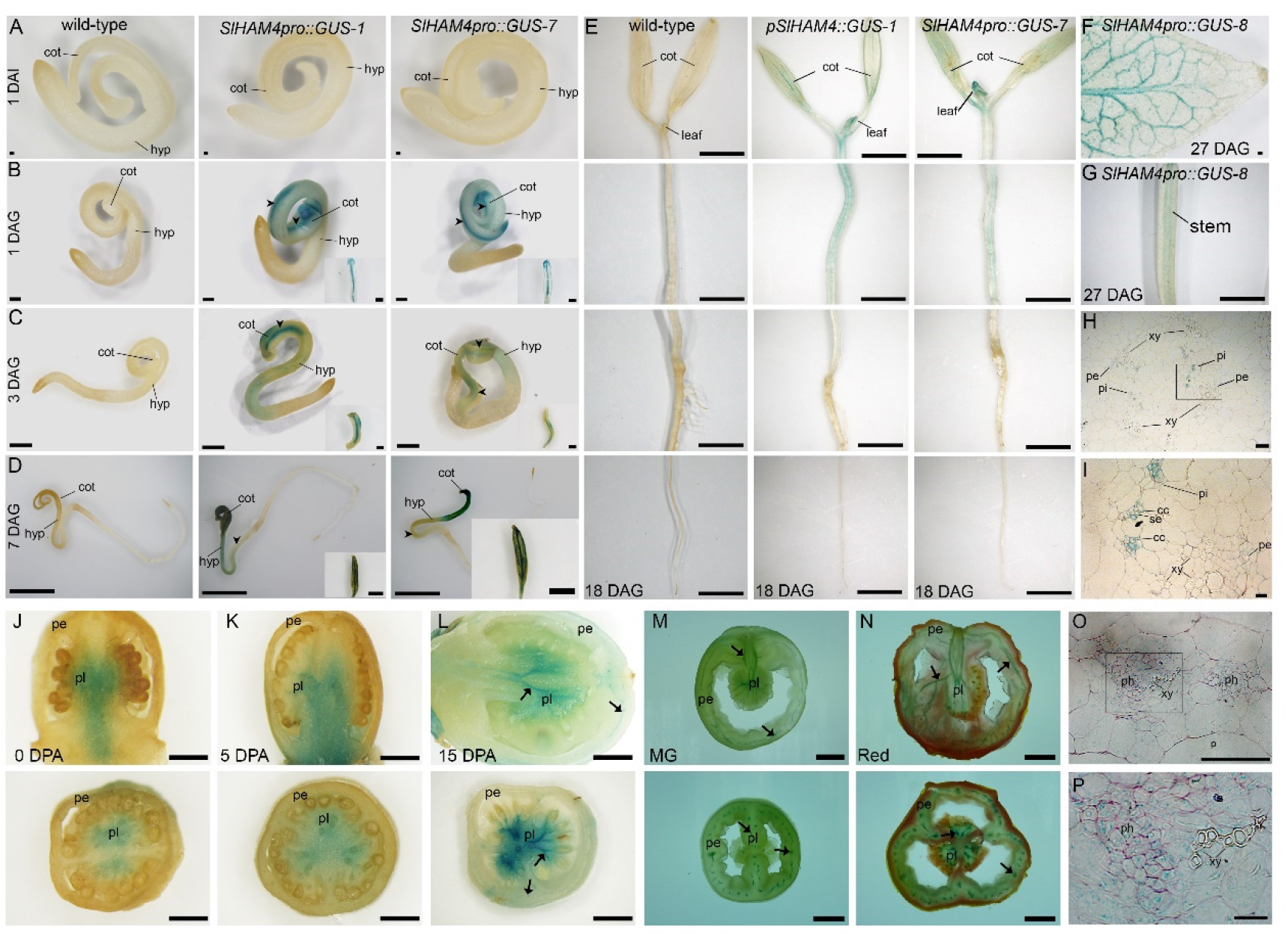
Tissue specific expression of *SlHAM4*. (A to O) Histochemical staining for GUS activity in M82 (wild-type) and transgenic tomato plants expressing GUS driven by *SlHAM4* native promoter (*SlHAM4pro::GUS*). GUS activity was visualized using the chromogenic substrate X-Gluc after ethanol clearing. (A) Mature embryos extracted from seeds one day after imbibition (DAI). (B to E) Representative whole seedlings at indicated days after germination (DAG). Insets in B to D show isolated cotyledons (adaxial side). (F) Adaxial side of the first leaf distal half. (G) Seedling stem. Scale bars = 200 µm (A and F), 500 µm (B), 1 mm (C), 2 mm (D, E and G). Inset scale bars = 500 µm (B and C), 1 mm (D). cot, cotyledon; hyp, hypocotyl. (J to O) Manual longitudinal (top panel) and cross (bottom panel) sections of anthesis ovary (J) and fruits at indicated developmental stage (K-N). DPA, days post anthesis. Arrows indicate GUS staining in representative vasculature. Scale bars 1 mm (J-L), 1 cm (M-O). (H, I, O and P) Histological cross sections of the *SlHAM4pro::*GUS-8 tissues stained with Ruthenium red. (H) Seedling stem shown in (G). (O) Pericarp vasculature of mature green fruit shown in (M). Scale bar = 100 µm. (I and P) Higher magnification of the histological sections outlined in (H) and (O) by a black box. Scale bar = 20 µm. p, parenchyma; ph, phloem; pi, internal phloem; pe, external phloem; xy, xylem; se, sieve element cell; cc, companion cell.

### Transcriptomic profiling of *slham4^CRΔ4^* ovaries

We utilized RNA sequencing (RNA-seq) to explore the impact of *SlHAM4* absence on the ovary transcriptome and gain insights into the molecular basis underlying tomato fruit catfacing. Comparative transcriptome analysis was conducted on -2DPA (stage 18) ovaries from wild-type, heterozygous (*slham4^CRΔ4(-/+)^*), and homozygous (*slham4^CRΔ4^*) M82 plants. For each genotype, three biological replicates of stage 18 ovaries were collected for RNA-seq library construction. Each library yielded approximately 16.5-19.5 million clean sequences, mapped to the tomato genome cDNA ITAG 2.5 (Table S1A). Notably, the 2^nd^ biological replicate of the *slham4^CRΔ4(-/+)^* group (*slham4^CRΔ4(-/+)^*-2) showed deviation (Fig. S6) and thus was excluded from subsequent analysis. Principal component analysis (PCA) confirmed the reproducibility among biological replicates, highlighting sample differences (Fig. 6A). Applying a fold-change cutoff of greater than 2 and a false discovery rate (FDR) below 0.01 as significance thresholds, we identified 475 differentially expressed genes (DEGs) in *slham4^CRΔ4^* ovaries compared to wild type. Among these, 142 were upregulated (up) and 333 were downregulated (down) (Fig. 6B and Table S1B). In comparison, the *slham4^CRΔ4(-/+)^* ovaries exhibited 140 up and 88 down DEGs (Fig. 6B and Table S1C). Of identified DEGs, 74 up and 39 down were unique to *slham4^CRΔ4(-/+)^* ovaries, while 77 up and 283 down were unique to *slham4^CRΔ4^* ovaries (Fig. 6C). Given that fruit catfacing was exclusively observed in *slham4^CRΔ4^* fruits, we considered the unique DEGs in their ovaries as potential candidate DEGs (cDEGs) contributing to catfacing. Moreover, 15 overlapping DEGs (1 up and 14 down; Table S1D), exhibiting similar trends and more pronounced fold changes in *slham4^CRΔ4^* compared to *slham4^CRΔ4(-/+)^*, were identified. The differential expression between *slham4^CRΔ4(-/+)^*and *slham4^CRΔ4^* mutants suggests that these DEGs might be regulated by SlHAM4 and could contribute to fruit catfacing. Consequently, these overlapping DEGs were also defined as cDEGs, bringing their total count to 375 (297 down, 78 up; Table S1E).

**Figure 6.**
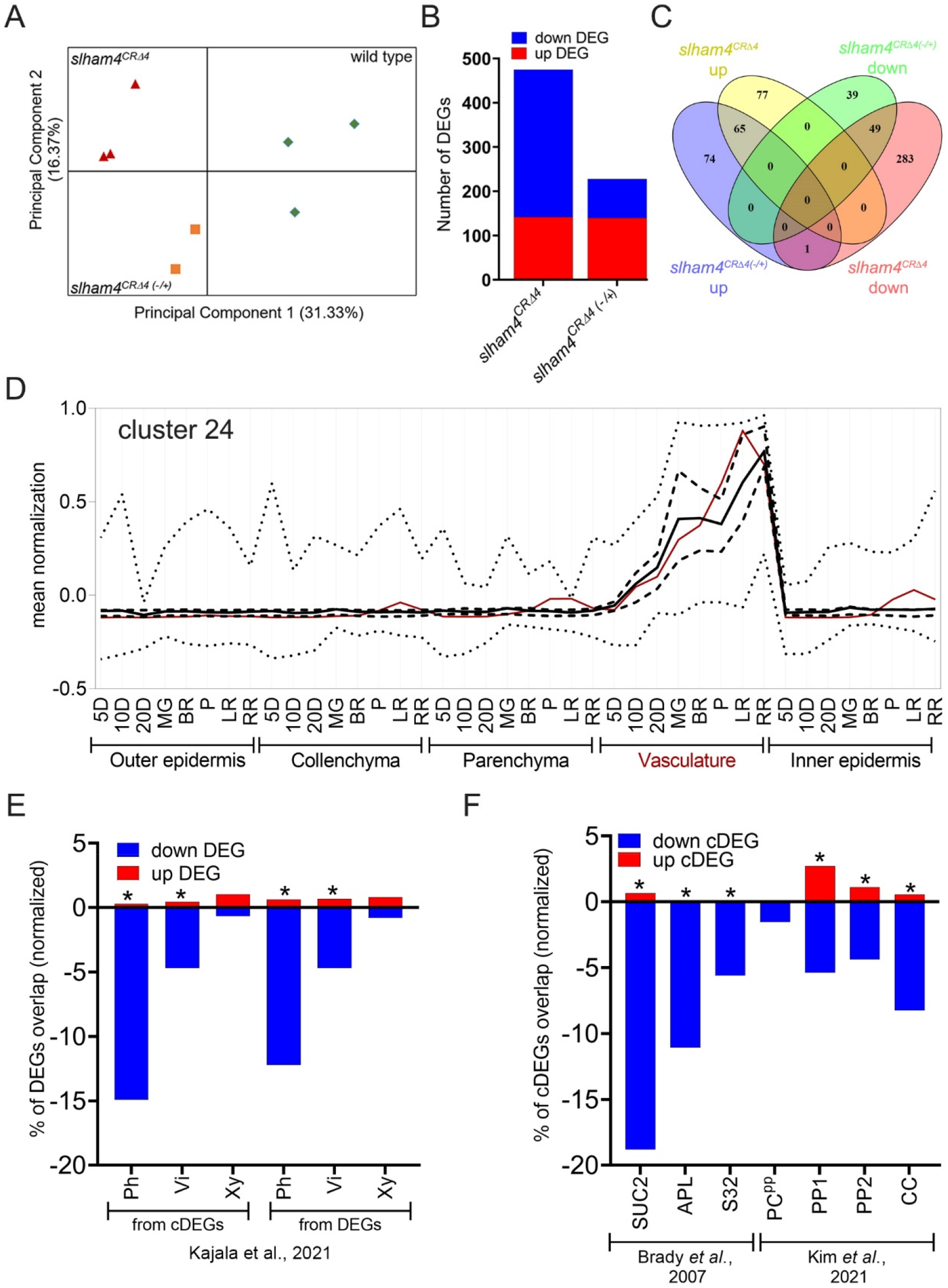
Global gene expression changes in *slham4^CRΔ4^* stage 18 ovaries. (A) Principal components analysis of all expressed genes showing three distinct groups. (B) Total number of DEGs in *slham4^CRΔ4^* mutant and heterozygous (*slham4^CRΔ4(-/+)^*) ovaries compared to wild-type. (C) Venn diagram displaying specific and overlapping DEGs between *slham4^CRΔ4^* and *slham4^CRΔ4(-/+)^* data sets. (D) Expression profile of cluster 24 cDEGs in fruit pericarp tissues based on TEA database data (Fernandez-Pozo *et al*., 2017). Cluster-wide average expression is plotted with solid lines, first and third quartiles by dashed lines, and maximum and minimum by dotted lines. The *SlHAM4* specific profile is plotted with red solid line. (E and F) Overlap between DEGs and cDEGs (E) or Arabidopsis orthologs of cDEGs (F) with indicated published vascular datasets. To compensate for dataset size, the numbers of up and down DEGs and cDEGs were normalized relative to the total number of genes in the largest group dataset (Supplementary Table S1F). Asterisks indicate statistical significance of overlap as calculated by http://nemates.org/MA/progs/overlap_stats.cgi. Ph, phloem; Vi, vascular initials; Xy, xylem.

### The expression of phloem-associated genes is altered in *slham4^CRΔ4^* ovaries

Utilizing gene expression data from the Tomato Expression Atlas (TEA) (Fernandez-Pozo *et al*., 2017), we performed a cluster analysis of the cDEGs alongside *SlHAM4*.

This analysis aimed to identify cDEGs that are co-expressed with *SlHAM4* in the same cells, a prerequisite for their transcriptional regulation by the SlHAM4 protein. We found that 51 down and 7 up cDEGs co-cluster with *SlHAM4* in cluster 24, which is mainly expressed in the vasculature of developing and especially ripening fruit pericarp (Fig. 6D and Table S1E). An additional 92 cDEGs exhibited a pericarp vasculature predominant expression (Fig. S7, clusters 2, 22, 23, 26, 27, 38), bringing the total number of vascular-associated cDEGs to 150, of which 91% (136/150) were downregulated in the *slham4^CRΔ4^* ovaries. To further explore the impact of *SlHAM4* loss- of-function on ovary vasculature transcriptome, we compared *slham4^CRΔ4^* DEGs with tomato genes found to be translated in root phloem, vascular initials, or xylem (Kajala *et al*., 2021) (Table S1F). This comparison revealed that 43% (25/58) of DEGs in cluster 24 are translated in the tomato root phloem, with negligible representation in root vascular initials and xylem (Table S1E). Moreover, 12.8% of the DEGs and 15.2% of the cDEGs were identical to root phloem genes. By contrast, only 3.2% of the DEGs and 1.3% of the cDEGs were identical to root vascular initials and root xylem genes, respectively (Fig. 6E and Table S1E). To identify phloem cell types whose transcriptome was affected due to *slham4* loss-of-function, we first matched each cDEG with its closest Arabidopsis ortholog, then checked for its presence in published Arabidopsis gene lists associated with specific phloem cells in roots (Brady *et al*., 2007) and leaves (Kim *et al*., 2021). This bioinformatic approach identified 36 and 35 cDEGs homologous to Arabidopsis root and leaf phloem-associated genes, respectively, mostly downregulated. Notably, the overlap was greatest with CC-specific genes in both root (SUC2, 10/36) and leaf (CC, 26/35) (Fig. 6F). Intriguingly, the leaf CC-specific gene list includes the gene code for HAM4 (AT4G36710), the Arabidopsis homolog of SlHAM4, and the Arabidopsis orthologs of 17.24% (10/58) of cluster 24 genes (Table S1E).

We annotated the tomato orthologs of key Arabidopsis genes involved in phloem development among the downregulated cDEGs. These include the tomato orthologs of the MYB transcription factor ALTERED PHLOEM DEVELOPMENT (APL; Solyc12g017370), crucial for protophloem SEs differentiation (Bonke *et al*., 2003); NAC-DEPENDENT EXONUCLEASE 1 (NEN1; Solyc03g115780), which promotes SE nuclear degradation and is transcriptionally regulated by the APL targets NAC45 and NAC86 (Furuta *et al*., 2014); LATERAL ROOT DEVELOPMENT 3 (LRD3; Solyc05g052910), involved in phloem development and function (Ingram *et al*., 2011); PHLOEM INTERCALATED WITH XYLEM (PXY; Solyc05g051640), a kinase critical for cambium cell divisions (Fisher and Turner, 2007); XYLEM INTERMIXED WITH PHLOEM 1 (XIP1; Solyc04g077010), regulating SE cell morphology (Tabata *et al*., 2014); VND7-INTERACTING 2 (VNI2; Solyc03g097650), a NAC transcription factor involved in phloem specification (Yamaguchi *et al*., 2010); ATP-BINDING CASSETTE G14 (ABCG14; Solyc08g075430), required for phloem development (Le Hir *et al*., 2013); and the CC-specific HSP20-like chaperone (AT5G54660/NPCC8, Solyc07g064020) (Zhang *et al*., 2008).

In addition, we annotated the tomato orthologs of Arabidopsis genes essential for phloem functions among downregulated cDEGs. These include the phloem-localized sulfate transporter SULTR1,3 (Solyc12g056930) (Yoshimoto *et al*., 2003); Phloem Protein 2 (PP2) and related lectins (Solyc02g069060, Solyc02g069020, Solyc02g069030, Solyc03g121300, Solyc00g048510, Solyc10g078600) that are expressed in CCs and thought to be involved in the long-distance movement of RNAs and defense (Pallas and Gómez, 2013); five orthologs of Thioredoxin, including three of Thioredoxin h (TRXh) (Solyc05g006830, Solyc05g006850, Solyc05g006870), which has been identified as a major protein in the phloem exudates of various monocots and dicots (Ishiwatari *et al*., 1995; Schobert *et al*., 1998; Carella *et al*., 2016) and was detected in rice leaf companion cells (Ishiwatari *et al*., 2000); CLAVATA1 (Solyc04g081590), which was shown to be expressed in root and leaf CCs and regulates lateral root outgrowth under N-deficient conditions (Araya et al., 2014; Brady et al., 2007); two orthologs of MYB-RELATED PROTEIN 2 (MYR2, Solyc10g085620, Solyc10g083340), linked to nitrogen uptake and assimilation; FT-INTERACTING PROTEIN 1 (Solyc03g077920), interacting with FLOWERING LOCUS T (FT) in the CCs to mediate its phloem export into SEs (Liu *et al*., 2012); EARLY FLOWERING MYB PROTEIN (EFM; Solyc01g108300), a root and leaf CCs expressed MYB transcription factor that is involved in negative regulation of flowering (Yan *et al*., 2014) and a tomato ortholog of SODIUM POTASSIUM ROOT DEFECTIVE 1 (NaKR1; Solyc10g085910), a phloem metal binding protein that is expressed in CCs and necessary for phloem function including the long-distance movement of FT (Zhu *et al*., 2016).

### The morphology of phloem is altered in *slham4^CRΔ4^*

Our results indicate that *SlHAM4* is expressed in phloem-associated cells including CCs (Fig. 5H, I, O and P), and its absence appears to modify their transcriptome (Fig. 6). This prompt us to further investigate *SlHAM4* specific functions in the phloem. Firstly, we assessed whether specific expression of *SlHAM4* in CCs could rectify the abnormal phenotypes observed in the *slham4^CRΔ4^* mutant. To that end, we transformed *slham4^CRΔ4^* plants with a binary construct expressing SlHAM4-GFP fusion protein driven by the Arabidopsis *SUCROSE-PROTON SYMPORTER 2* (*AtSUC2*) promoter (Stadler and Sauer, 1996), shown to be CC-specific also in tomato (Spiegelman *et al*., 2015) (Fig. S3F). This transformation led to the regeneration of seventeen *AtSUC2pro::SlHAM4 slham4^CRΔ4^* T0 transgenic plants (Fig. S3G). However, these plants still exhibited sepal browning during flower senescence and fruit catfacing (Fig. 4C), suggesting that *SlHAM4* specific expression in CCs is insufficient to rescue the *slham4^CRΔ4^* phenotypes. In Arabidopsis, the phloem mutants *xylem intermixed with phloem1* (*xip1*) and *phloem intercalated with xylem* (*pxy*) are known to accumulate anthocyanin in their shoots (Bryan *et al*., 2012; Fisher and Turner, 2007). In our *slham4^CR^* mutants, anthocyanin over-accumulation was observed in young leaves and sepals (Fig. 2D and F), coinciding with downregulation of the orthologs of XIP1 and PXY (Table S1E). To investigate whether anthocyanin over-accumulation is linked to abnormal phloem development, we conducted a comparative anatomical analysis of the vasculature in the leaf rachis and flower pedicel of *slham4^CRΔ4^* and wild-type tomato plants. In the wild type, vascular bundles are typically U-shaped in the leaf rachis (Fig. 7A and B) and ring-shaped in the flower pedicel (Fig. 7E and F). These vascular tissues comprise external phloem bundles adjacent to the cortex, surrounded by xylem vessels, and internal phloem bundles near the pith periphery (Fig. 7B, C, F and G). These phloem bundles contain SEs, which are relatively small compared to surrounding cells, interspersed with smaller companion cells (insets Fig. 7C and G). Histological analysis of the *slham4^CRΔ4^*leaf rachis and flower pedicel revealed no significant alterations in the organization or composition of these vascular bundles (Fig. 7C, D, G and H), and their counts remained comparable to the M82 (Fig. 7I, O, L and R). However, a notable reduction in the area of external phloem bundles was observed in the mutant, along with a decreased cell count in these bundles (Fig. 7J, K, P and Q). We also observed reductions in the mutant internal phloem bundle area and respective cell count, but these differences were not statistically significant (Fig. 7M, N, S and T).

**Figure 7.**
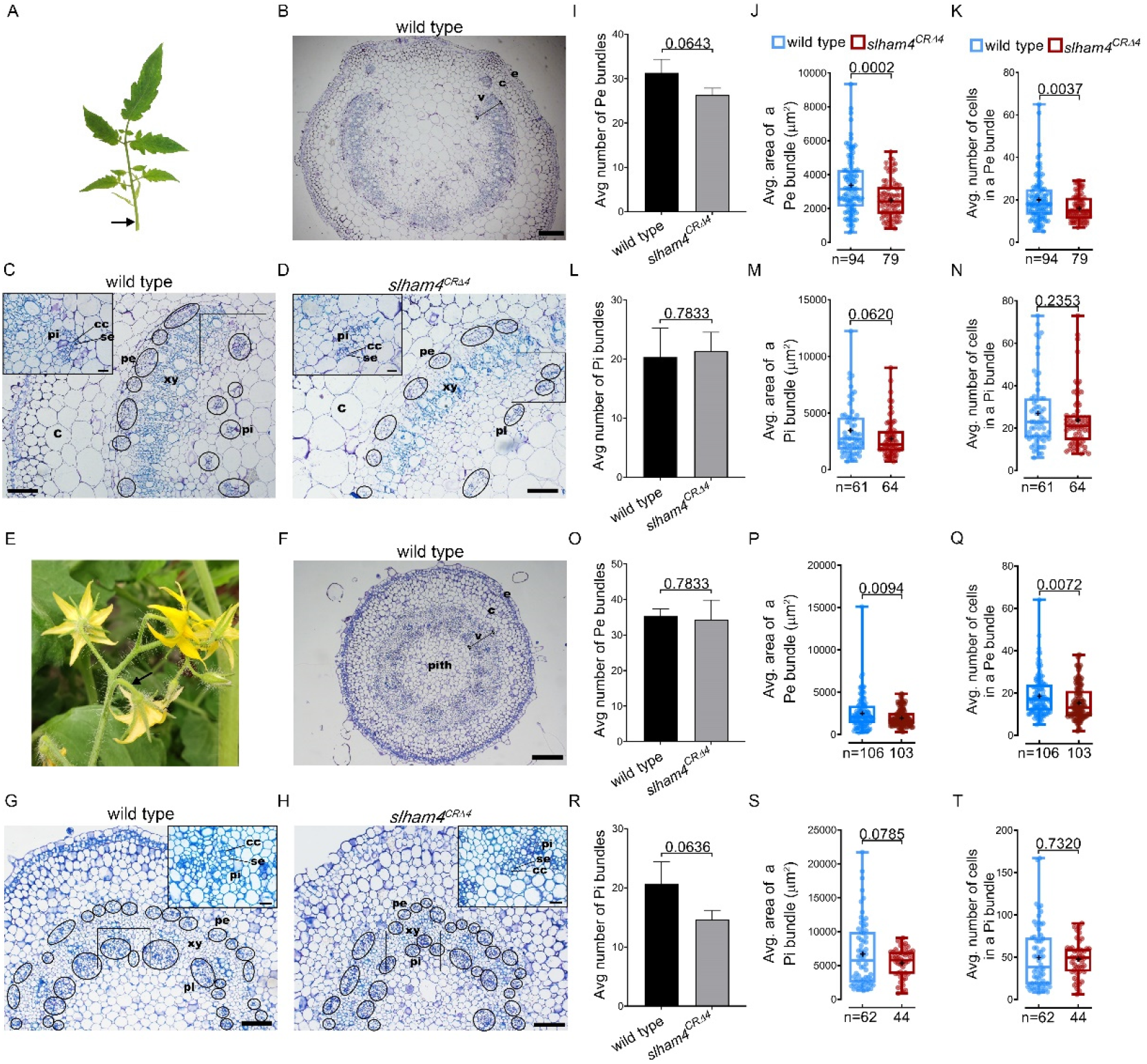
Histological comparison of wild-type and *slham4^CRΔ4^* leaf rachis and flower pedicel phloem bundles. Representative pictures of fully expanded leaf (A) and anthesis flower (E). Arrows indicate the rachis (A) and pedicel (E) regions analyzed. Representative Toluidine blue stained plastic-embedded cross sections of leaf rachis (B to D) and flower pedicel (F to H). Cross section of whole rachis (B) and pedicel (F), the vasculature tissue (v) is delimited by a black line. Representative higher magnification cross sections of rachis (C and D) and pedicel (G and H). The external (pe) and internal (pi) phloem bundles are circled in black and insets show higher magnification of the phloem bundles outlined by a black box. The small angular cells that are almost filled with blue contents are companion cells (CCs). Scale bars = 200 µm (B), 100 µm (C, D and F), 50 µm (G and H), 20 µm (inset C and D), 10 µm (inset G and H). e, epidermis; c, cortex; xy, xylem; se, sieve element cell. Measurements of rachis (I-N) and pedicel (O-T) average phloem bundle number (I, L, O and R), area (J, M, P and S) and cell count (K, N, Q and T). Error bars indicate ±SD over 2 biological replicates (I, L, O and R). Box plots center line, median; box limits, upper and lower quartiles; whiskers, min and max values; points, individual values; + = mean; n = number of phloem bundles. The *P* values as determined by Student’s t-test are shown.

## Discussion

This study presents a comprehensive functional analysis of *SlHAM4* in tomato, uncovering its roles in development and phloem function. Our findings suggest that SlHAM4 is involved in gene regulation within the phloem system.

Public expression data and GUS reporter assays demonstrated that *SlHAM4* is predominantly expressed in the vasculature of various organs. The plant vasculature is composed of dead xylem, which conducts water and minerals, and living phloem, which transports vital nutrients such as sugars, amino acids, and hormones throughout the organism (Hardtke, 2023). Histochemical GUS staining revealed that within the vasculature bundle, *SlHAM4* expression is confined to phloem tissues, contrasting with an absence of expression in the nearby xylem. This phloem-specific expression is supported by the detection of *SlHAM4* mRNA within the phloem translatome of tomato seedling roots (Kajala *et al*., 2021) and is consistent with the expression of Arabidopsis *HAM4/SCL15* detected by GUS staining in *SCL15pro::GUS* seedlings (Gao *et al*., 2015). The lack of severe developmental defects in the phloem of *slham4^CRΔ4^* and the nearly wild-type phenotype of the *ham4/scl15* Arabidopsis mutant (Gao *et al*., 2015), and the abundance of *HAM4* in the phloem of mature organs such as ripening fruits in tomato and mature roots in Arabidopsis (Gao *et al*., 2015), suggest that SlHAM4 is involved in phloem functionality and may play a minor or redundant role in phloem development.

Phloem tissue primarily comprises three main cell types: the enucleate SEs that form the sieve tube, the adjacent CCs which support the SEs through connecting plasmodesmata and phloem parenchyma cells (Hardtke, 2023). Within the phloem tissue, GUS staining detected the expression of *SlHAM4* in phloem associated cells including CCs. These results are in concordance with observations in Arabidopsis, where the SCL15/HAM4-GFP fusion protein was localized to phloem-specialized cells, including CCs (Gao *et al*., 2015), and single-cell transcriptomics identified HAM4 mRNA specifically in the CCs of leaves (Kim *et al*., 2021). Moreover, our transcriptomic analysis reveals that the absence of SlHAM4 activity leads to reduced expression of several genes known to be expressed in CCs. This likely influences the protein composition of the phloem, thereby affecting its functionality. A significant reduction in the expression of TRXh genes, known for their roles in maintaining the proper function of various target proteins by ensuring correct disulfide bonding (Gelhaye *et al*., 2004), in *slham4^CRΔ4^*, exemplifies the potential impact of SlHAM4 on phloem protein content. Members of the TRXh family are known for their expression in CCs and abundance in the phloem (Reichheld *et al*., 2002). The decrease in TRXh levels within the phloem could potentially compromise target protein functionality by allowing the oxidation of redox-sensitive cysteines. Nevertheless, specific expression of *SlHAM4* in CCs failed to rectify the phenotypic abnormalities observed in *slham4^CRΔ4^*mutants, implying that the roles of SlHAM4 extends beyond CCs to other cells of the phloem. Supporting this, the SCL15 protein was also identified in the phloem parenchyma cells of leaf petioles (Gao *et al*., 2015) and in early vascular precursor cells, such as those in the provascular or procambial tissues (Zhou *et al*., 2015b). These findings collectively suggest a broader regulatory role for SlHAM4 in various aspects of phloem function.

In tomato, phloem-mediated transport of photosynthates and essential nutrients is vital for meeting the metabolic requirements of the developing fruit, a major sink organ, thereby supporting its growth (Ho, 1996). We observed a marked size reduction in *slham4^CR^* fruits compared to those of the M82 cultivar, pointing to inhibited growth in the former. This and the specific *SlHAM4* expression in the phloem of developing fruits raise the possibility that compromised phloem function in *slham4^CR^* mutant fruits may impede the transport of photosynthates and essential nutrients, adversely affecting their normal growth. Furthermore, our study revealed that the absence of functional *SlHAM4* lead to the development of small ruptures in the pericarp of an expanding set fruit, ultimately causing significant scarring in mature mutant fruits. This phenotype closely resembles the Catface syndrome in tomatoes, a disorder characterized by large, irregular scars on the fruit’s blossom end (Peet, 2009; Srinivasulu *et al*., 2020). Research suggests that fruit catfacing originates from the incomplete closure of carpels at the base of the style in rapidly growing fertilized ovaries. This defect may result from the over proliferation of ovary tissues, particularly the locules and style, during flower development. Accordingly, brief cold stress or gibberellin treatment during flowering promotes ovary tissue proliferation such as style fasciation and increased locule number, which is associated with increased fruit catfacing (Sawhney, 1982). Recent research has demonstrated a link between cold stress and the disruption of the WUS- CLV3 feedback loop, which leads to stem cell proliferation in the tomato flower meristem (Wu *et al*., 2022). This disruption provides a possible explanation for cold-induced tomato fruit catfacing. However, the *slham4^CR^* mutant flowers display normal-looking pistils at anthesis, indicating that catfacing in these mutants is not due to over proliferation of ovary tissues. Notably, in certain cold-treated tomato varieties, fruit catfacing was not associated with ovary tissue proliferation. It was suggested that cold temperatures might inhibit the formation or transport of growth substances critical for normal cell division near the style’s base, preventing complete carpel closure (Knavel and Mohr, 1969). This raise the possibility that suboptimal phloem function in *slham4^CRΔ4^* anthesis ovaries fails to support the rapid growth of the setting fruit pericarp leading to the formation of small ruptures that eventually enlarge into scars, although the specific reason for scar formation at particular locations remains unclear.

In summary, our results suggest that tomato *HAM4* is a phloem-associated gene that is necessary for proper development, as well as for phloem integrity, in part by modulating the expression of phloem genes, particularly those expressed in CCs. Further research are expected to elucidate which CC genes are transcriptionally regulated by SlHAM4, and their specific contribution to phloem functions.

## Materials and methods

### Plant material and growth conditions

Germinated tomato cv. M82 seedlings and regenerated T0 plants were grown in a growth chamber at 24 ^ο^C under 50-70 μmol m^-2^ s^-1^ photosynthetic photon flux density with 16/8 hr light/dark period. Subsequently, one-month-old seedlings and acclimated regenerated plants were transplanted into four-liter pots containing a tuff-peat mix soil. These plants were grown under greenhouse conditions, with temperatures maintained between 25 °C and 30 °C (day/night). Crosses were made in emasculated flowers by hand-pollination.

### Plasmids constructions

For pBI101-SlHAM4pro::GUS binary plasmid, a 4000 bp sequence upstream of the *SlHAM4* start codon was PCR amplified from the M82 tomato genomic DNA using the primer pair SlHAM4pro-BamHI-F1 and SlHAM4pro-XmaI-R that contained *BamH*I and *Sma*I flanking restriction sites. Following *BamH*I and *Sma*I digestion the amplified fragment was cloned into the corresponding sites of the binary plasmid pBI101 just upstream of *GUS*. For the pART27-35S::SlHAM4 binary plasmid, a 1611 bp encoding *SlHAM4* was PCR amplified from an anthesis ovary cDNA with the primer pair SlHAM4-SalI-Fwd and SlHAM4-HindIII-Rev (primer sequences can be found in Table S2). The amplified fragment was digested with *Sal*I and *Hind*III and subsequently cloned into *Xho*I and *Hind*III sites of the pART27-35S binary vector downstream of the *35S* promoter. The pDGB3-Omega1-SlHAM4pro::SlHAM4-GFP::TNOS and pDGB3- Omega1-AtSUC2pro::SlHAM4-GFP::TNOS binary plasmids were constructed using the GoldenBraid cloning system. *AtSUC2* promoter and *GFP* (with Gly-Gly-Ser linker) were PCR amplified from the *AtSUC2pro*::*GFP* tomato line genomic DNA (Spiegelman *et al*., 2015). The 4Kb SlHAM4pro, *AtSUC2* promoter and *GFP* PCR amplicons were individually cloned into the GoldenBraid entry vector (level 0), pUPD2 (Sarrion-Perdigones *et al*., 2011). The expression cassette was precisely generated in level 1 binary plasmid pDGB3-Alpha1 by assembling *SlHAM4pro/AtSUC2pro*, *SlHAM4* and *GFP* with additional *NOS* terminator (*TNOS*). The constructed expression cassette was further integrated with plant kanamycin resistance cassette pNOS::nptII::TNOS in the level 2 binary plasmid pDGB3-Omega1 to generate the final binary plasmids. All binary constructs were validated by sequencing. Primer sequences used for plasmid constructions are listed in Table S2.

### CRISPR/Cas9-mediated mutagenesis of *SlHAM4*

First, two gene-specific gRNAs targeting the coding region of *SlHAM4* were designed (gRNA sequences are provided in Table S2) and each was incorporated *in silico* into sgRNA . Then, a construct containing the two sgRNAs in tandem each driven by the synthetic Arabidopsis U6 promoter delimited by 5’-*Mlu*I and 3’-*Hind*III was artificially synthesized (GeneWiz, USA) and cloned into pUC57 to generate pUC57-U6::sgRNA1- U6::sgRNA2. The pUC57-U6::sgRNA1-U6::sgRNA2 plasmid was digested with *Mlu*I and *Hind*III restriction enzymes, and the released U6::sgRNA1-U6::sgRNA2 fragment was ligated into the corresponding sites of pRCS binary vector, which also contained a plant codon-optimized version of Cas9 driven by the 35S promoter (Damodharan *et al*., 2018). To detect the CRISPR/Cas9-induced mutations in *SlHAM4*, the genomic DNA was extracted from each transgenic T0 plant and screened by PCR for the presence of the *35S::Cas9* transgene with the primer pair Cas9-Fwd and Cas9-Rev. Then, the *SlHAM4* targeted sequences in the transgenic T0 plants were PCR amplified with primer pair SlHAM4-Cas9-valid-F and SlHAM4-Cas9 valid-R and sequenced to identify indels in them. The T0 plants carrying mutation were backcrossed to M82 wild type, and the resulting F1 progeny plants were genotyped for the absence of the *35S::Cas9* transgene and presence of indels. The identified non-transgenic F1 CRISPR mutants were selfed, and homozygous F2 CRISPR mutants were genotyped by sequencing. Detection of homozygous *slham4^CRΔ4^* and *slham4^CRΔ3^* mutants was further done by PCR with the primer pair CR-SlHAM4delta4detect-fwd and SlHAM4-Cas9 valid-R, which amplify only the *SlHAM4* wild-type allele. Primer sequences are listed in in Table S2.

### Tomato transformations

Binary constructs were transformed into tomato cv. M82 (pBI101-SlHAM4pro::GUS, pRCS-2xU6syn::SlHAM4, pART27-35S::SlHAM4) or *slham4^CRΔ4^* (pDGB3-Omega1- SlHAM4pro::SlHAM4-GFP, pDGB3-Omega1-AtSUC2pro::SlHAM4) plants by co-cultivation of 12 day-old cotyledons with *Agrobacterium* strain GV3101 as described previously (Damodharan *et al*., 2016).

### Histochemical GUS staining

For GUS staining, *SlHAM4pro::GUS* tomato transgenic tissues were submerged in GUS staining buffer [100 mM sodium phosphate buffer (pH 7.0), 2 mM potassium ferricyanide, 2 mM potassium ferrocyanide, 0.1% Triton X-100, 10 mM EDTA and 1 mM X-Gluc] following by 1 hr of vacuum infiltration with six intermittent stops and overnight incubation in 37 °C. Three washes with 70% ethanol (v/v) followed by incubation at 37 °C for 8 hr after each wash dechlorophylized the stained tissues, which were then visualized or used for histology.

### Histology

GUS stained and unstained tissues were fixed in PFA solution as described in Hendelman et al., (2016). Microtome-cut sections of 4-μm thick were mounted on microscopic slides and stained with 0.1% (*w/v*) Toluidine blue for 1 min (flower pedicel and leaf rachis) or 0.1% (*w/v*) Safranin for 1 hr followed by 0.1% (*w/v*) Fast Green for 30 s (ovary and fruit). For GUS stained tissues microtome-cut sections of 10-μm thick were mounted on microscopic slides and further stained with 0.05% (*w/v*) Ruthenium red for 1 min. Slides were examined and photographed under bright field using an Olympus DP73 microscope equipped with a digital camera.

### Phloem bundles analysis

M82 and *slham4^CRΔ4^* leaf rachis and flower pedicel external and internal phloem bundles were analysed in Toluidine blue stained plastic-embedded cross sections of three independent biological replicates per each tissue. A total of 3 and 2 parts per single leaf rachis and flower pedicel biological replicate, respectively, were analysed manually, or in the case of phloem bundle area, by using ImageJ 1.53e software (http://imagej.nih.gov/ij) as described in Figure S8.

### Transcriptome analysis by RNA-seq

Total RNA was extracted from isolated ovaries using Bio-Tri RNA reagent (Bio-Lab, Israel) as described by the manufacturer’s protocol. For RNA-seq, three biological replicates of wild type, *slham4^CRΔ4^* and *slham4^CRΔ4(-/+)^* stage 18 (-2 DPA) isolated ovaries were used. Each replicate contained 10 ovaries from 8 independent plants. RNA-seq libraries (Illumina Truseq RNA) preparation and pair-end sequencing were performed at MACROGEN-EUROPE (Macrogen, The Netherlands). Differential expression analysis was done at the ARO bioinformatic unit. Briefly, raw reads underwent a filtering and cleaning procedure. The SortMeRNA tool was used to remove rRNA sequences. The FASTX Toolkit (version 0.0.13.2) was then employed to trim read-end nucleotides with quality scores below 30 using the FASTQ Quality Trimmer, and to discard reads with less than 70% base pairs having a quality score of 30 or higher using the FASTQ Quality Filter. Reads were mapped to the tomato coding sequences (ITAG2.4 release provided by the International Tomato Annotation Group) using Bowtie2. Transcript quantification was performed with the Expectation-Maximization method (RSEM), utilizing the align_and_estimate_abundance.pl script from the Trinity software package (https://github.com/trinityrnaseq/trinityrnaseq/wiki). Principal Component Analysis (PCA) was conducted using R Bioconductor. Differential expression analysis was carried out with the edgeR R package, considering genes with a false discovery rate (FDR) below 0.01 and at least a 2-fold change as differentially expressed.

### Quantitative PCR analyses

Total RNA was extracted from indicated tissue using Bio-Tri RNA reagent (Bio-Lab) as described by the manufacturer’s protocol. Two micrograms of total RNA were treated with DNase I, followed by 40 cycles of PCR to ensure the absence of genomic DNA in the samples. First-strand cDNA was then synthesized from one microgram of total RNA using a Maxima first-strand cDNA synthesis kit (Thermo Scientific, Lithuania) according to the manufacturer’s instructions. qPCR was performed on a StepOnePlus system (Thermo Scientific), and the results were analyzed with StepOne software version 2.2.2 (Thermo Scientific). Relative expression levels were normalized using SlTIP41 as a reference gene and calculated using the comparative delta delta Ct (ΔΔCt) method. The primers for qPCR are listed in Table S2.

### Bioinformatic Analysis

The Tomato Expression Atlas database (TEA; http://tea.solgenomics.net/) gene expression levels (average RPM) of *SlHAM4* and indicated DEGs in the fruit pericarp cell types across stages (5D to RR) were used for K-means clustering, that was done using Morpheus website (https://software.broadinstituteorg/morpheus) with number of row clusters = 40. The mean normalized value for each DEG per sample (fruit pericarp cell type at a specific stage) was calculated using the formula (X - Xav) / (Xmax - Xmin), where X is the average RPM value of a specific DEG, and Xav, Xmin, and Xmax are the mean, minimum, and maximum average RPM values of that DEG across all samples, respectively. The Arabidopsis orthologs of identified tomato DEGs were determined using the Best-Hits-and-Inparalogs (BHIF) method at https://bioinformatics.psb.ugent.be/plaza/versions/plaza_v4_5_dicots/download.

## Acknowledgements

We would like to thank Prof. Shmuel Wolf for providing the seeds of the *AtSUC2::GFP* tomato line. We also extend our gratitude to Dr. Amir Sherman and Dr. Ada Rozen for their assistance in genotyping the *slham4^CR^* mutants, and to Dr. Adi Faigenboim for her help in analyzing the RNA-seq results.

## Author contributions

A.P.V. O.G. S.C. characterized the *slham4^CR^* mutants. A.P.V. generated and characterized the *35S:SlHAM4* transgenic plants. S.K.G. preformed the RNAseq. J.K. generated and characterized the *SlHAM4pro::GUS*, *SlHAM4pro::SlHAM4-GFP* and *AtSUC2pro::SlHAM4* transgenic plants, analyzed wild-type and *slham4^CRΔ4^* phloem bundles and assisted in manuscript writing. T.A. Conceptualization, planned the experiments, supervised the work, annotated the RNA-seq data and wrote the manuscript.

## Supplementary data

**Figure S1.** Expression patterns of *SlHAM4* according to public databases.

**Figure S2.** Variation of the catface phenotype in *slham4^CRΔ4^* fruits.

**Figure S3.** Construction of complementation constructs and validation of transgenic plants.

**Figure S4.** Analysis of T0 *SlHAM4* GUS reporter plants.

**Figure S5.** Histological analysis of the *SlHAM4pro::GUS-8* cotyledon vasculature region.

**Figure S6.** Heatmap of hierarchically clustered Pearson correlation matrix for gene expression in wild-type and *SlHAM4* mutant samples.

**Figure S7.** Expression profiles of indicated cluster cDEGs in fruit pericarp tissues based on TEA database data.

**Figure S8**. Analysis of phloem bundles number, area and cell number per bundle.

**Table S1.** RNA-seq mapping statistics and lists of DEGs between stage 18 ovaries of wild type, *slham4^CRΔ4(-/+)^* and *slham4^CRΔ4^*.

**Table S2.** Primers and gRNAs used in this study.

## Data availability

The RNA-seq data is available from the SRA database under the accession number PRJNA682014. The candidate DEGs and *SlHAM4* expression profiles in pericarp cell types tissue across the stages (5DPA to RR) of tomato fruit were retrieved from the Tomato Expression Atlas database (http://tea.solgenomics.net/)(Shinozaki *et al*., 2018).

